# AMaNITA: an end-to-end workflow for native tRNA nanopore sequencing data analysis

**DOI:** 10.64898/2026.06.16.732588

**Authors:** Xanthi-Lida Katopodi, Leszek P Pryszcz, Laia Llovera, Alexane Ollivier, Luca Cozzuto, Julia Ponomarenko, Eva Maria Novoa

**Author notes:** Correspondence to: Eva Maria Novoa.

## Abstract

Transfer RNA (tRNA) molecules serve as essential adapters during protein translation. While direct RNA sequencing (DRS) via Oxford Nanopore Technologies has emerged as a powerful platform for systematic tRNAome profiling, we currently lack a simple and robust statistical framework for nanopore tRNA data analyses. Here, we address this gap by developing AMaNITA (Abundance, Modifications, and Nanopore Intensity Toolbox Application), an end-to-end bioinformatic workflow that enables simplified, robust, and scalable analyses of nanopore native tRNA sequencing datasets. AMaNITA streamlines the entire analytical trajectory: from upstream processing (basecalling, mapping, filtering, batch effect correction) to downstream assessment of differential tRNA abundance and modification stoichiometry. The workflow generates an interactive HTML report for data exploration and analysis, allowing the user to download the source data files and resulting plots. AMaNITA can be executed using Singularity from the command line, without requiring installation of dependencies.

## INTRODUCTION

Transfer RNAs (tRNAs) are highly modified, small non-coding RNAs that play an essential role in protein translation. The chemical modifications present in tRNAs are critical for their stability, and are dynamically regulated in response to environmental stimuli, cell cycle progression and tumorigenesis (1, 2). Similarly, tRNA abundances are frequently dysregulated upon environmental exposures and across various human diseases (3, 4).

Traditionally, tRNA modifications have been quantified using mass-spectrometry (MS)-based methods (5–9). While MS is highly sensitive, it typically cannot identify the specific position or exact tRNA isoacceptor where a modification occurs. Conversely, tRNA abundance is typically measured using next-generation-sequencing (NGS) methods (10–15), which requires conversion of the tRNA molecule into cDNA, a process that “erases” tRNA modifications (16). Although specialized NGS-based methods have incorporated variations in their reverse transcription (RT) enzyme choice or optimized buffer conditions to capture information from a subset of RNA modifications (12, 17, 18) –typically in the form of systematic “errors” such mismatches or indels)– they fail to provide a complete view of the tRNAome.

To overcome these limitations, recent studies have employed the direct RNA sequencing (DRS) platform developed by Oxford Nanopore Technologies (ONT) to characterize the tRNAome (19–29). To capture tRNAs using ONT DRS, two critical adaptations compared to standard DRS protocols (30) were required. First, tRNA molecules were extended to ensure full-length tRNA molecules, which was achieved using a splint ligation approach (20, 22). This approach was later optimized into “Nano-tRNAseq” (20), a gel-free protocol compatible with lower input samples. Second, adjustment of MinKNOW configuration parameters was required to prevent the software from prematurely discarding short tRNA molecules, a step that was essential for accurate, length-independent quantification of tRNA abundances (20). This adaptation has now been incorporated in the more recent versions of MinKNOW, compatible with the latest SQK-RNA004 direct RNA sequencing chemistry. More recently, the subsequent development of tRNA-specific multiplexing barcodes and demuxing models (26, 31, 32) has further enabled the use of Nano-tRNAseq as a simple and cost-effective method for simultaneous quantification of tRNA abundances and modification dynamics.

Despite these advances, analyzing the expanded lexicon of the tRNAome remains a significant challenge. While over 170 different RNA modification types have been identified to date –the majority of which reside in tRNAs (2, 33, 34)–, modification-aware basecalling models have been primarily trained for RNA modifications present in mRNA (35). Consequently, nanopore-based tRNA studies have relied largely on comparative analyses of raw signal features (27) or basecalling error profiles (20, 21). Although recent works have explored the applicability of modification-aware models to tRNA (36), their interpretability is confounded by the cross-reactivity of the models with other RNA modification types present in tRNAs (23, 37).

Regardless of the detection method employed, a largely unaddressed hurdle is the robust statistical identification of tRNA-modified positions that significantly change in response to a specific treatment, or upon comparison with a knockout strain. Moreover, the field currently lacks an end-to-end solution that integrates efficient upstream processing –including basecalling, mapping, counting, filtering, quality control, and batch effect correction– with rigorous downstream differential tRNA modification and abundance analyses.

Here, we address this gap by developing AMaNITA (**A**bundance, **M**odifications, **a**nd **N**anopore **I**ntensity **T**oolbox **A**pplication), an end-to-end workflow that allows for a simplified, robust, and scalable bioinformatic analysis of nanopore tRNA sequencing datasets, going from raw sequencing data to a final HTML report containing differentially expressed and differentially modified tRNA positions between groups of interest (**Fig 1A**, see also **Fig S1**). AMaNITA comprises two main parts: i) *AMaNITA Preprocess*, in which the reads are basecalled, demultiplexed and mapped using a tRNA-specific NextFlow workflow (**Fig 1B**) and ii) *AMaNITA Main*: a modular pipeline that generates interactive HTML reports (**File S1**), which include quality control analysis (quality metrics, filtering, batch effect correction) as well as differential tRNA modification and tRNA abundance analyses (**Fig 1C**). To demonstrate the utility of AMaNITA, we apply the workflow to a panel of seven *Saccharomyces cerevisiae* (yeast) knockout (KO) strains, each harboring a targeted knockout of a specific RNA-modifying enzyme, as well as to wild type and *Mus musculus* (mouse) *Dnmt2* knockout samples. Our analyses demonstrate that AMaNITA does not only accurately recapitulate known modification-enzyme relationships, but also identifies technical noise and batch effects that can hinder the accurate detection of tRNAome variations. Beyond simple detection, this toolbox provides a modular framework that can be easily adapted to any species of interest, offering a standardised method to explore tRNA dynamics across diverse biological and cellular contexts.

**Figure 1.**
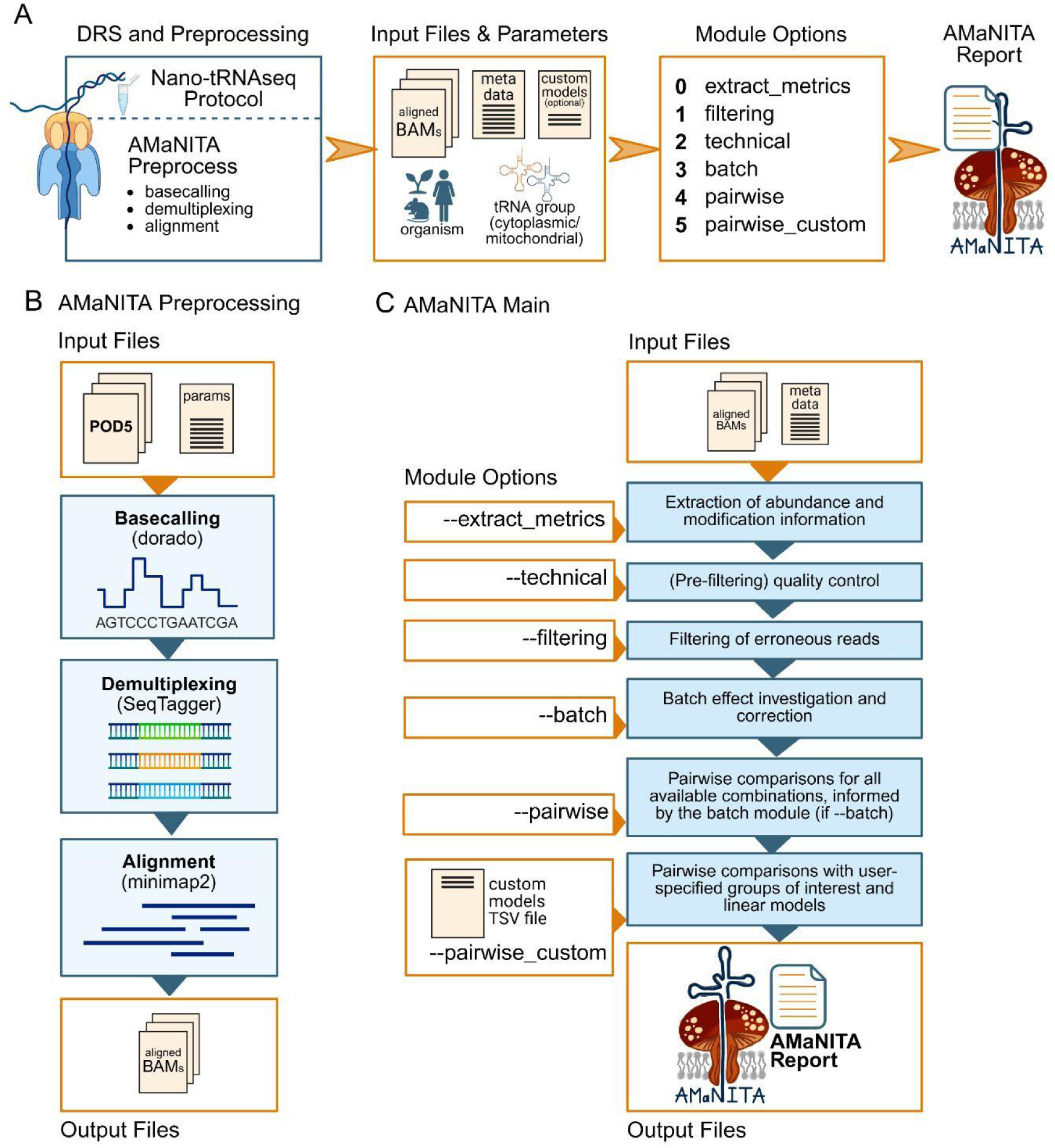
AMaNITA is an end-to-end workflow for nanopore tRNA sequencing data analysis. **(A)** Schematic representation of the steps performed to obtain processed reports with user-friendly plots starting from raw Nano-tRNAseq sequencing data, which requires two main steps: i) AMaNITA Preprocess: raw data processing by performing basecalling, demultiplexing (if barcoding is available), and alignment; ii) AMaNITA Main: analysis of tRNA abundances and modifications using the AMaNITA pipeline, an end-to-end pipeline that performs quality control (technical QC), filtering, batch correction, differential tRNA abundance analysis and differential tRNA modification analysis on the dataset of interest. **(B)** Schematic overview of AMaNITA Preprocess workflow, which consists in a tRNA-optimized NextFlow workflow that performs basecalling (dorado), demultiplexing (SeqTagger) and aligning (minimap2) tRNAs, utilizing the MasterOfPores version4 (MoP4) preprocess module with tRNA-specific parameters. **(C)** Schematic overview of AMaNITA Main workflow, which uses aligned BAM files as input and produces an AMaNITA Report and plots as output. This is performed by running 6 individual modules in AMaNITA Main, which include five core modules (extract_metrics, technical, filtering, batch, pairwise), and one optional module (pairwise_custom).

## RESULTS

### AMaNITA Preprocess: a Nextflow Workflow for Nano-tRNAseq Data Pre-processing

Compared to mRNA reads, which are typically mapped using ONT “default” recommended parameters, tRNA molecules require specific optimized mapping configurations and stringent filtering (20, 22, 38, 39). These adjustments are required to accommodate the high density of tRNA modifications, which induce increased basecalling errors (20); without such optimizations, a substantial proportion of tRNA reads would remain unmapped. To streamline these non-standard steps, we developed *AMaNITA Preprocess*, a tRNA-specific adaptation of the MasterOfPores (40, 41) Nextflow workflow. This pipeline is pre-configured with parameters optimized for Nano-tRNAseq reads, ensuring technical replicability and computational reproducibility. The workflow includes three main steps: (i) *<u>basecalling</u>* using dorado (https://github.com/nanoporetech/dorado), (ii) *<u>demultiplexing</u>* via SeqTagger (31), and (iii) sensitive *<u>alignment</u>* to a curated, reduced tRNA reference library using minimap2 (42), employing parameters specifically tuned for error-prone tRNA reads (see *Methods*) (**Fig 1B**).

### AMaNITA: an end-to-end solution for tRNA nanopore sequencing data analysis

The AMaNITA pipeline consists of five main modules (**Fig. 1A**): (i) the *<u>extract_metrics</u>* module extracts tRNA counts and modification information from BAM files, generating tables which are required as input for other AMaNITA modules; (ii) the *<u>filtering</u>* module performs identification and filtering of low quality, misaligned, and erroneous reads; (iii) the *<u>technical</u>* module performs quality control of the dataset; (iv) the *<u>batch</u>* module performs batch effect investigation and, upon the identification of batch effects that contribute to the variance, it corrects for them; and (v) the *<u>pairwise</u>*module, which performs pairwise comparisons between groups of interest that have been specified in a user-provided metadata file, performing both differential tRNA abundance (expression) and differential modification analysis. The pairwise module is also available in a *<u>pairwise_custom</u>*version, which allows the user to define which groups to compare, linear models to adopt, or comparisons to make (e.g. one vs many groups, or many vs many) (**Fig 1C**).

AMaNITA is designed to compare two biological conditions (e.g. knock-out (KO) versus wild type (WT), treated versus control, disease versus normal, etc). The results generated by the individual modules are then automatically concatenated into a single HTML report with tables and interactive plots, as well as in the form of standalone files and figures that can be used for downstream analyses and publication purposes. In addition, the user can select which modules to turn on/off, with each combination of settings producing a separate report (**Fig 1A**). We should note that a minimum of three replicates per condition are recommended (43); however, AMaNITA can also work with fewer replicate designs. The AMaNITA code is containerized, and can be easily executed from the command line via Singularity. The code is publicly available in GitHub and is distributed with a GPL-3.0 license (see *Code Availability* section).

### Quality control in AMaNITA

Quality control in AMaNITA occurs in three modules: the *<u>filtering</u>*, the *<u>technical</u>*, and the *<u>batch</u>* modules. We illustrate the analyses and plots that can be obtained using these three modules in a panel of 16 *S. cerevisiae* Nano-tRNAseq samples, consisting of wild type and 7 knockout strains (**Fig 2A**), in biological duplicates, which were sequenced as barcoded samples in different flowcells (**Table S1** and **S2**).

**Figure 2.**
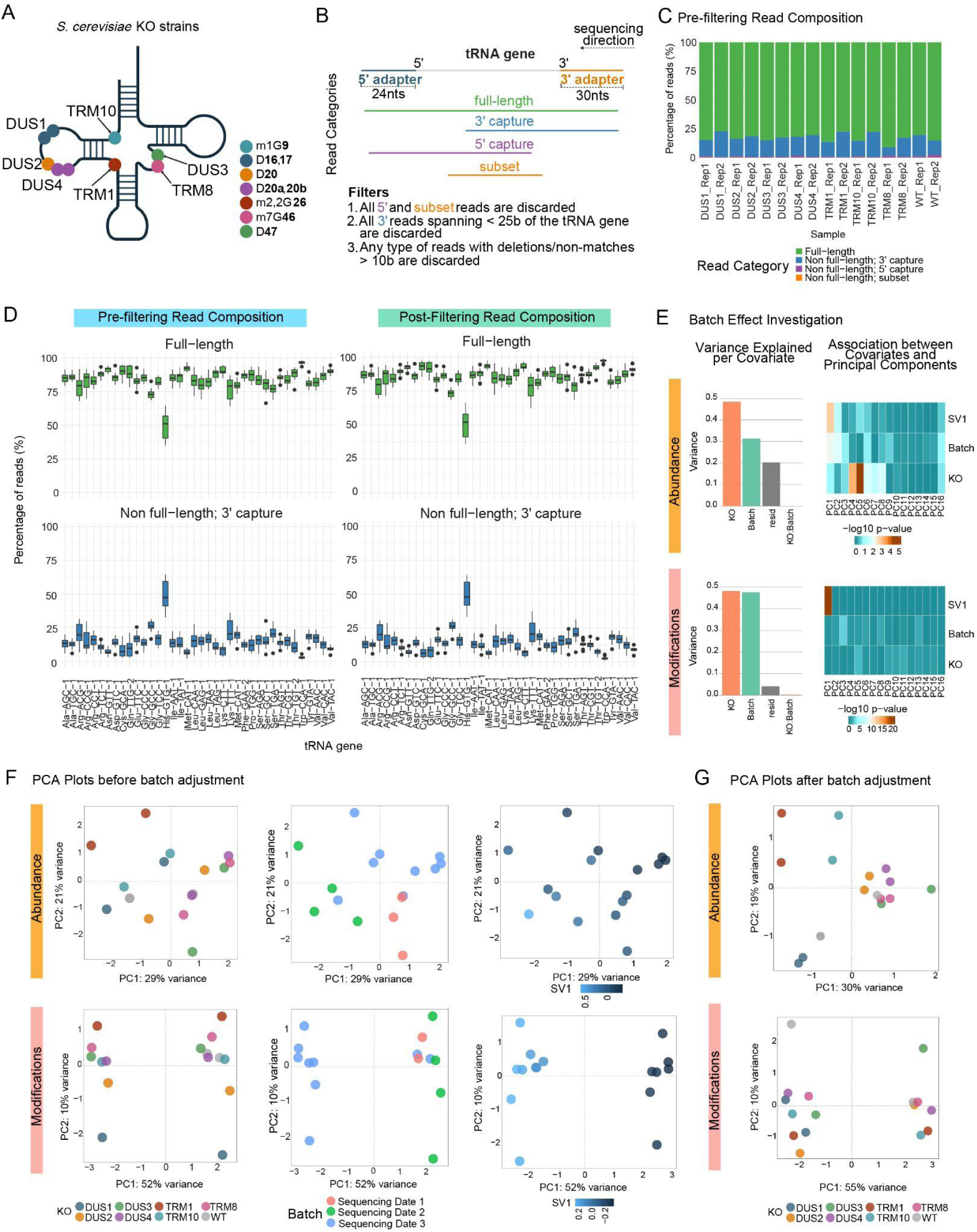
Quality control analysis of *S. cerevisiae* wild type and tRNA modification knockout (KO) strains using AMaNITA. **(A)** Schematic representation of the positions and modifications placed by the RNA modifying enzyme that is knocked-out in the strains used in this work: TRM10 (m^1^G9), DUS1 (D18, D19), DUS2 (D20), DUS4 (D20a, D20b), TRM1 (m22G26), TRM8 (m7G46), DUS3 (D47). **(B)** AMaNITA’s filtering module classifies the reads into four different categories: i) full-length reads, ii) 3’ capture reads, iii) 5’ capture reads and iv) subset reads. Only full-length reads and 3’ capture reads are used by downstream analysis modules. See also **Tables S3** and **S4** for tRNA counts before and after filtering, respectively. **(C)** Proportion of the four read categories defined by AMaNITA across the different yeast KO Nano-tRNAseq samples, before the filtering step. **(D)** Boxplots depicting the proportion of full-length (top) and 3’ capture (bottom) categories of reads, before (left) and after (right) the filtering step, generated by AMaNITA’s *filtering* module. **(F)** Abundance (top) and modification (bottom) exploratory PCA plots, coloured by biological group (left), sequencing run -suspected batch effect-(middle), and identified surrogate variable (SV1) (right). **(G)** Abundance (top) and modification (bottom) exploration of the biological and technical variance of the data as identified by Principal Variance Component Analysis (PVCA, left) and by plotting the p-value of the association of principal components (PCs) with covariates of interest (right). **(G)** PCA plots of tRNA abundance (left) and modification (right) data, after batch correction by adjusting for the identified SV1. See also **Tables S3** and **S4** for tRNA abundances (counts) per tRNA reference, before (S3) and after (S4) filtering..

The *<u>filtering</u>* module identifies potentially misaligned reads, and excludes them from downstream analyses. Misaligned reads are identified based on three assumptions: (i) direct RNA sequencing proceeds in the 3’-to-5’ direction, therefore, reads should always map from the 3’ end, and should not initiate at the middle of the gene body; (ii) *bona fide* reads should contain the 3’ Nano-tRNAseq adapter, as sequencing requires the presence of a polyA tail –which is included in the 3’ adapter–; and (iii) tRNA reads should span a substantial portion of the tRNA gene, since very short fragments are unlikely captured by MinKNOW, and therefore are more likely to represent misaligned reads. Based on this criteria, mapped reads are classified into four categories: (i) full-length reads, comprising the 3’ adapter, tRNA gene, and 5’ adapter, (ii) 3’ capture reads, whose alignment begins at the 3’ adapter and spans portion of the tRNA genes but the read is truncated before reaching the 5’ adapter, (iii) 5’ capture reads, spanning part of the tRNA gene and 5’ adapter, and (iv) subset reads, which only span part of the tRNA gene without capturing 3’ or 5’ adapters (**Fig 2B**). AMaNITA considers that reads falling into the latter two categories are likely to correspond to spurious reads or erroneous alignments, as they do not meet the three rules described above; consequently, they are filtered out. In addition, 3’ capture reads covering less than 25 bases of the tRNA are also excluded, since we have consistently identified these reads as misaligned. In the case of the yeast KO panel, we observe 75-90% full-length reads across samples (**Fig 2C**). We should note that AMaNITA additionally offers the option of working only with full-length reads or only with truncated (3’ capture) reads, which might be of interest for certain experimental settings and designs.

The *<u>technical</u>* module performs quality control of the dataset, providing sample- and tRNA-specific technical metrics. This module allows identifying samples that may be problematic in the dataset (**Fig 2C**), or examining whether tRNAs are evenly represented across the samples included in the dataset. For example, in the *S. cerevisiae* dataset (**Table S1**), the technical module identifies tRNA-His(GTG) to be enriched in 3’ capture reads compared to other tRNAs, both before and after filtering (**Fig 2D**, see also **Fig S2**). This behaviour is expected, as it is caused by the G-1 modification that occurs at the -1 position of tRNA-His(GTG), which does not allow proper ligation of the 5’ adapter, leading to an apparent loss of ∼15 bases of tRNA-His(GTG), in agreement with previous works (20).

The *<u>batch</u>* module identifies batch effects potentially contributing to the variance of the data, assesses their significance, and if identified, suggests them as covariates for the linear models in downstream analyses (**Fig 2E**). The batch module includes exploratory data analysis visualizations in the form of Principal Component Analysis (PCA) biplots (**Fig 2F**) and Hierarchical Clustering plots (**Fig S3A**). These plots allow users to visually examine whether their datasets segregate following their “conditions” of interest (e.g. treated and untreated), or whether additional variables (e.g., library preparation date, flowcell) might be contributing in the form of batch effects. If potential batch effects are provided by the user in the form of metadata (see for an example **Table S2**), AMaNITA carries out Principal Variance Component Analysis (PVCA) (*44*), which identifies the percentage of variance that is explained by non-biological or technical variables (**Fig 2E**). In addition, AMaNITA implements Surrogate Variable Analysis (SVA) (45, 46) to agnostically extract sources of technical variation in the form of surrogate variables from the data itself (47). Upon the identification of all sources of technical variation, AMaNITA follows a pre-determined set of rules for selecting the variables that should be accounted and corrected for (see *Methods*). The batch module then provides batch-corrected PCA plots (**Fig 2G**), which uncover how the data clusters based on the biological “conditions” of interest rather than technical noise, and then informs the pairwise module to account for those sources of technical variation in the linear models while performing differential analyses. We should note that while AMaNITA guides model selection, users still can visualize and obtain both batch-adjusted and non-batch-adjusted results.

In the *S. cerevisiae* dataset, batch effect investigation using PVCA revealed a dominant batch effect correlated with the sequencing run (sequencing date), which was found to explain 30% and 45% of the variance in the tRNA abundance and modifications data, respectively (**Fig 2E**). In agreement with this, the agnostic batch effect analysis using SVA revealed one significant surrogate variable that significantly correlated with PC1 after dimensionally reducing both the abundance and modifications data (**Fig 2E**). These results illustrate that batch effects can be present in direct RNA sequencing runs, and can be corrected for using batch-effect-correcting methods (**Fig 2F**, see also *Methods*).

It is important to note that the *batch* module should be used with caution when the datasets are not adequately balanced, i.e. when the variable causing the suspected batch effect (for example, sequencing date) is confounded with the biological effect of interest (**Fig S3B**). In these cases, AMaNITA will print a “warning” to the user (**Fig S4**), in which the user is warned that correcting for batch effects in an unbalanced experiment can lead to artefacts in the prediction of biological signal.

### Differential tRNA abundance and modification analyses using AMaNITA

Differential tRNA abundance (expression) analysis in AMaNITA is carried out using the DESeq2 package (48). DESeq2 uses a generalized linear model (GLM)-based approach, and allows for complex experimental designs that include multiple biological or technical covariates, as well as multiple replicate handling and multi-group comparisons. AMaNITA then visualizes the results in the form of volcano plots and hierarchically-clustered tRNA abundance heatmaps (**Fig S5 and S6**).

AMaNITA uses two different metrics as a proxy for differential tRNA modifications: i) differential basecalling errors, which have been previously used for RNA modification detection (20, 21, 29, 49), and ii) differential basecalling quality score (baseQ) signal (50). Identification of differentially modified positions using basecalling errors is performed using edgeR (51), adopting the differential methylation mode previously used to assess differential DNA methylation states, e.g. for a bisulfite-treatment assay (WGBS or RRBS) (52). Conversely, differential modification analysis via baseQ is performed using the Kolmogorov-Smirnov (K-S) test (53), which allows comparison of baseQ distributions between groups. While the K-S test is a computationally efficient approach for handling baseQ scores, it does not inherently support complex linear modeling or batch effect correction. Therefore, this module is best suited for balanced designs where technical variability caused by batch effects is minimized.

In the *S. cerevisiae* dataset, we observed a strong batch effect confounded with the biological signal, due to its unbalanced experimental design (**Fig S3**). In those cases, AMaNITA has built-in warnings to alert users of potential bias in the results (**Fig S4**), since attempting to adjust for the batch effect in an unbalanced experimental design can lead to incorrect biological conclusions (**Fig S7** middle panels). On the contrary, analyzing the data from the same KOs but with a properly balanced experimental design (see **Table S1**) yielded the expected results (**Fig S7** right panels).

We then examined the performance of AMaNITA in more complex mammalian tRNAomes. Previous works have reported that N5-methylcytosine (m^5^C) modifications are among the most challenging RNA modification types to detect using ONT direct RNA sequencing (49, 54, 55). For this reason, we sought to examine whether AMaNITA would correctly identify m^5^C modifications by comparing *M. musculus Dnmt2* knockout (KO) samples to their wild type (WT) counterparts, which are expected to lack m^5^C at position 38 exclusively in tRNA^Asp^ (56) (**Fig 3A**). To this end, we sequenced tRNAs from both WT and *Dnmt2* KO mouse testis samples, using both RNA002 and RNA004 chemistries. AMaNITA revealed a batch effect in the RNA002 dataset, correlated with the flowcell and sequencing date, whereas no batch effect was identified for the RNA004 counterpart (**Fig 3B**). Differential modification analysis revealed that the expected loss of the m^5^C modification at position 38 could be correctly identified only after batch adjustment in the RNA002 dataset. Similarly, AMaNITA correctly identified the modified position in the RNA004 dataset (**Fig 3C**). We should note that the differential modification was observed at a shifted position (i.e., position 39 (+1) for the RNA002 dataset, and position 37 (-1) for the RNA004 dataset) (**Fig 3C**), in agreement with previous works reporting a shifted basecalling “error” signal of m^5^C modifications in DRS datasets (49). Notably, the “shifted” position varied across the two chemistries, most likely due to the use of distinct *dorado* versions of canonical basecalling models, which were not equivalent for RNA002 and RNA004 chemistries.

**Figure 3.**
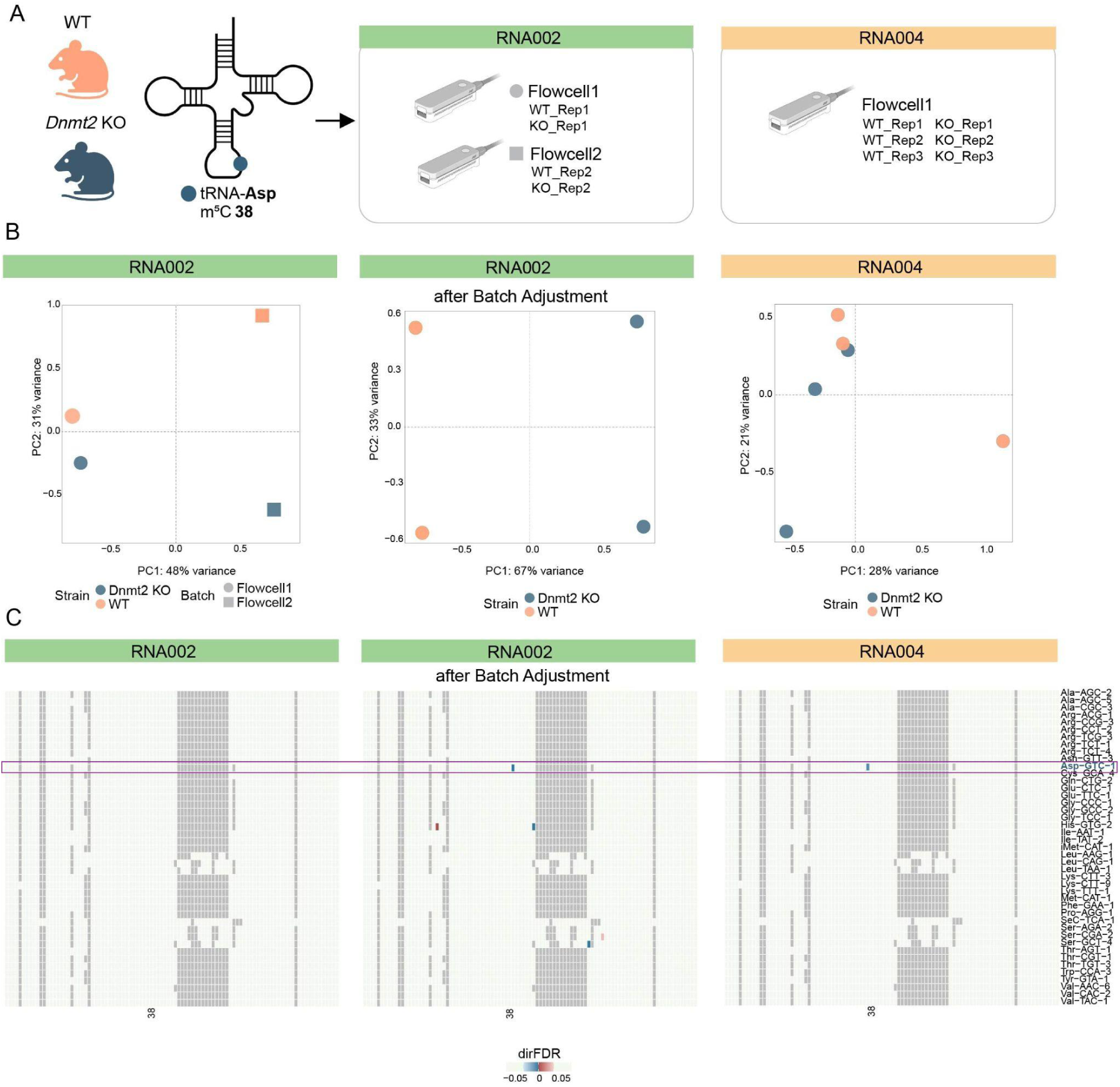
Example of differential tRNA modification analysis performed by AMaNITA, showcased on *M. musculus* Dnmt2 KO strain (compared to wild type). **A)** Schematic representation of the experimental design: small RNAs from wild type (WT) and *Dnmt2* KO testis tissue samples were sequenced using RNA002 or RNA004 chemistries, in biological duplicates or triplicates. For RNA002, samples were sequenced in two separate barcoded flowcells. For RNA004, samples were sequenced in a single flowcell, with barcodes. The expected specificity of *M.musculus Dnmt2* enzyme is also depicted (tRNA-Asp m^5^C38). **B)** PCA plots of tRNA modification (using basecalling “errors” as proxy for modification information) coloured by the biological group (WT or KO). The shapes (square or circle) represent the identified batch effect. **C)** PCA plot of tRNA modifications (using basecalling “errors” as proxy for modification information) coloured by the biological group for the mouse KO dataset, sequenced with RNA004 chemistry. **D)** Positional heatmaps depicting the results of the differential modification analyses of the mouse *Dnmt2* KO versus WT. In the left panel, results are shown for the RNA002 dataset without adjusting for batch effect. In the middle panel, results for the RNA002 dataset adjusting for batch effect are shown. In the right panel, results for the RNA004 dataset without adjusting for batch effect are shown. Each row is a tRNA (collapsed reference, see also *Methods*) and each column represents a canonical position of the tRNA. The heatmaps are colored based on their significance (assessed by directional false discovery rate (FDR), see also *Methods*). Positions in red correspond to significant increase in tRNA modification frequency, whereas positions in blue correspond to significant decrease in tRNA modification frequency. Positions in white represent non-significant changes in tRNA modification frequency.

## DISCUSSION

In recent years, it has become possible to sequence native tRNA reads, providing information-rich snapshots of the tRNAome at an unprecedented resolution (19–29). A major unsolved challenge, however, has been the lack of a simple and robust bioinformatic pipeline to facilitate the analysis native tRNA populations. Notably, their short length, dense modification content, and high basecalling error rates make that bioinformatic pipelines for tRNA analysis will be fundamentally different from those used for mRNAs, justifying the need for dedicated mapping, filtering, and modification detection strategies (20, 22, 25, 27, 32, 57). Moreover, existing approaches for tRNA analysis are typically fragmented, often focus only on signal or basecalling errors, and do not offer comprehensive solutions that integrate preprocessing, QC, batch correction and statistical testing in a single framework.

AMaNITA addresses this gap by providing an end-to-end workflow for nanopore tRNA sequencing data analysis, from raw reads to differential tRNA abundance and modification results, which can be visualized in an interactive HTML report (see **File S1** as an example). Firstly, raw data is processed using a NextFlow workflow, which performs basecalling, demultiplexing, and mapping including tRNA-optimized sensitive alignment mapping parameters. Then, AMaNITA is used to perform quality control, filtering likely misaligned and spurious reads. It then informs and corrects for potential batch effects. With the filtered and batch-corrected datasets, it performs differential tRNA abundance and modification stoichiometry analyses.

To identify which tRNA nucleotides –and in which tRNA isoacceptor– are differentially modified, AMaNITA uses two metrics: differential basecalling errors (computed as the sum of mismatches, deletions and insertions at a given position (20, 21, 29)), and baseQ. Alternative approaches to detect tRNA modifications have consisted in the use of current signal differences (39) as well as in differential per-read modification probabilities, generated by *dorado* modification-aware models (23). However, their applicability of the latter will be limited to a subset of tRNA modifications that may show cross-reactivity with modification-aware basecalling models (23).

In this work, we demonstrate how AMaNITA’s statistical framework manages to assess the quality of a Nano-tRNAseq dataset, identify sources of technical variation, prioritize the most significant ones to be adjusted for, and accurately capture differential abundance and modification events. Notably, we demonstrate that RNA modification estimation can greatly suffer from batch effects in native tRNA sequencing runs, which had not been previously appreciated. We should note that direct RNA sequencing (DRS) has been traditionally assumed to be exempt from batch effects, due to their simplified library preparation steps and lack of PCR amplification. Here we demonstrate that these batch effects exist, and can contribute to noise in the detection of modifications (false positives) as well as lack of statistical significance (false negatives) (**Fig S7**).

We should note, however, that AMaNITA still presents limitations. Firstly, its detection of modifications relies on pairwise comparisons; it is not capable of *de novo* detection of tRNA modifications, and cannot identify the chemical identity of the site that is predicted to be differentially modified. For this reason, orthogonal methods may still be needed if the ultimate goal is chemical identification. In this regard, the possibility of integrating LC-MS/MS into Nano-tRNAseq pipelines can offer a promising solution to address this limitation (58). Secondly, AMaNITA relies on mapping the tRNAs to reduced tRNA reference sets rather than employing the full tRNAome reference set, which might mask tRNA isoacceptor-specific tRNA modification differences. While some efforts have been made towards achieving tRNA isoacceptor resolution (25), this is still an unsolved computational problem, and the current proposed solutions are far from being applicable to real biological datasets, especially in the case of complex tRNAomes found in higher eukaryotes. Thirdly, we should note that depending on the RNA modification type, AMaNITA may not predict RNA modifications with single nucleotide resolution, but rather, it will identify a “region” (subset of consecutive nucleotides) as differentially modified. This is a well-known limitation of using basecalling “errors” as a proxy for identifying modifications, where some RNA modification types typically yield single-nucleotide position errors (e.g., pseudouridine (Y) (20), m^5^C (49), N6-methyladenosine (m^6^A) (50)) but others produce broader error signals (e.g. 2’-O-methylations (49), N1-methyladenosine (m^1^A)(59), dihydrouridine (D), N2, N2-dimethylguanosine (m22G)), typically spanning 3-4 nucleotides (**Fig S7** and previous works (20, 49, 50, 59)).

Overall, AMaNITA provides an end-to-end solution for Nano-tRNAseq data analysis. We envision that it will accelerate our understanding of tRNA dynamics upon stress, across development, and in disease. Future work could extend AMaNITA to incorporate modification-aware models, as well as to integrate it with orthogonal methods for tRNA modification detection, such as LC-MS/MS.

## MATERIALS AND METHODS

### Mouse husbandry

Animal experimentation was carried out in compliance with EU Directive 86/609/EEC and Recommendation 2007/526/EC regarding the protection of animals used for experimental and other scientific purposes, enacted under Spanish law 1201/2005, and also approved by the institutional ethics committee. Wild type mice used in this study were of C57BL/6J strain background, obtained from Charles Rivers (strain #000664). *Dnmt2* knockout mice (B6;129-*Trdmt1^tm1Bes^*/J) were obtained from Jax Laboratories (strain #006240), and were back-crossed with C57BL/6J strain mice for 6 generations. All mice were raised on a defined control diet (Special Diets Services, RM1 (P), 801151). Mice were housed in cages at a temperature of 22–24°C, had access to food and water *ad libitum* and were maintained on a 12:12 hour light-dark artificial lighting cycle, with lights off at 19:00.

### Tissue collection and small RNA extraction from mouse testis

Mice were euthanized with CO_2_ in compliance with ethics approval. Testis tissues were dissected, snap-frozen and stored at -80°C until tissue RNA extraction. Small pieces of tissue sample (50–100 mg) were homogenized in 1 mL of TRIzol using a Polytron homogenizer (Tekmar TISSUEMIZER). 200 μL of chloroform was added, after vortexing the samples briefly and a 3 min incubation at RT, samples were centrifuged at 12, 000 × g for 15 min at 4°C. The aqueous phase was transferred to a new tube, and an equal volume of 70% ethanol was added. After vortexing thoroughly, 700 μL of the mixture was transferred to a RNeasy Mini spin column (Qiagen, 74104) the samples were centrifuged at 12, 000 × g for 30 sec at RT, the flow-through which contains the small RNAs fraction was kept. The small RNAs were purified using the RNeasy MinElute Kit (Qiagen, 74204). 0.65X of 100% ethanol was added to the flow-through, and the mixture was transferred to a RNeasy MinElute spin column. After centrifugation at 12, 000 × g for 30 sec at RT, the flow-through was discarded. Columns were washed with 700 μL of RWT buffer, centrifuged at 12, 000 × g for 30 sec and then 500 μL of RPE buffer was added to the column and samples were centrifuged at 12, 000 × g for 30 sec. Finally, columns were washed with 500 μL of 80% ethanol and centrifuged at 12, 000 × g for 2 min to dry the membrane. The column which contains the small RNAs was placed into a 1.5 mL collection tube. Samples were eluted by adding 17 μL of RNase-free water directly onto the column and centrifuged at 12, 000 × g for 1 min. RNA concentration was quantified using Qubit, and the quality of the separation was verified by using the Agilent TapeStation.

### S. cerevisiae strains and culturing

*S. cerevisiae* knockout strains (BY4741 MATa trm1::KAN, BY4741 MATa dus1::KAN, BY4741 MATa dus2::KAN BY4741, MATa dus3::KAN, BY4741 MATa dus4::KAN, BY4741 MATa trm8::KAN, BY4741 MATa trm10::KAN) were obtained from the Yeast Knockout Collection (Dharmacon) and were grown under standard conditions in YPD medium (1% yeast extract, 2% Bacto Peptone and 2% dextrose) at 30 °C. Starting from a 4mL O/N cultures, these were diluted to 0.0001 OD600 in 200 ml of YPD and grown overnight at 30 °C with shaking (250 r.p.m.). When cultures reached the mid-exponential growth phase (OD600 0.5), they were transferred into a pre-chilled 50-ml Falcon tube and centrifuged at 3, 000g for 5 minutes at 4 °C, followed by two washes with water. Pellets were snap-frozen at −80 °C.

### Total RNA extraction and short RNA separation

Snap-frozen yeast pellets were resuspended in 660 µl of TRIzol Reagent (Thermo Fisher Scientific, 15596018) with 340 µl of acid-washed and autoclaved 425–600-µm glass beads (Sigma-Aldrich, G8772). Cells were disrupted by vortexing at maximal speed for seven cycles of 15 seconds and chilling the samples on ice for 30 seconds between cycles. The samples were then incubated at room temperature for 5 minutes, and 200 µl of chloroform (Vidra Foc, C2432) was added. After briefly vortexing the suspension, samples were incubated for 5 minutes at room temperature and centrifuged at 14, 000g for 15 minutes at 4 °C. The upper aqueous phase was transferred to a new tube. To precipitate RNA, 1× volume of molecular-grade isopropanol and 1 µl of GlycoBlue co-precipitant (Thermo Fisher Scientific, AM9515) were added and mixed by inverting and incubated for 10 minutes at room temperature. The samples were centrifuged at 14, 000g for 15 minutes at 4 °C, and the pellet was then washed with ice-cold 70% ethanol. The pellet was resuspended in 20 µl nuclease-free water after air drying for 5 minutes at the benchtop. RNA purity was measured using a NanoDrop 1000 spectrophotometer. Then, 10 µg of each sample was treated with Turbo DNase (Thermo Fisher Scientific, AM2238) and subsequently cleaned up using a Zymo RNA Clean and Concentrator-5 kit (Zymo Research, R1016) following the manufacturers’ instructions to keep only the small RNA fraction (<200 nt). The short RNA fraction was eluted in nuclease-free water. RNA concentration was determined using Qubit Fluorometric Quantitation; RNA purity was measured with a NanoDrop 1000 spectrophotometer; and the RNA electropherogram was obtained using Agilent 4200 TapeStation RNA HS ScreenTape Assay.

### tRNA deacylation

An input of 10 uL of small RNAs was mixed to 90 uL of 100mM Tris-HCl pH 9. Samples were incubated 30 min at 37°C. Samples were then cleaned-up using Zymo RNA Clean and Concentrator-5 kit (Zymo, R1016) following the manufacturer’s instructions to retain RNAs ≥17 nt. RNA was eluted in 10 uL of RNase-free water. The RNA concentration was quantified by using Qubit Fluorometric Quantitation.

### Nanopore direct tRNA sequencing library preparation (Nano-tRNAseq)

Nanopore tRNA libraries were prepared as previously described (20) with minor modifications. Briefly, 500 ng of deacylated RNA was ligated to the NanotRNA-seq adapter overnight at 4°C. Then, custom barcoded RTA adapters were used in place of the standard ONT RTA to enable sample multiplexing. Barcoded RNAs were reverse transcribed by Maxima H Minus RT for 30 min at 60°C, pooled and purified using 1.8× AMPure RNAClean XP beads. Then, tRNA libraries were prepared using either the SQK-RNA002 or SQK-RNA004 kit following the manufacturer’s instructions and sequenced on MinION devices using FLO-MIN106 or FLO-004RA flowcells, respectively.

### AMaNITA Preprocess

AMaNITA Preprocess employs MasterOfPores version 4.1 (MoP4.1) (40, 41), configured with parameters optimized for Nano-tRNAseq reads. Basecalling was performed using dorado (https://github.com/nanoporetech/dorado) version 1.3.1, using the “fast” basecalling models (rna004_130bps_fast@v5.0.0 for RNA004 datasets and rna002_70bps_fast@v3 for RNA002 datasets). Demultiplexing was performed using SeqTagger version 1.1 (31), using tRNA-specific demuxing models from SeqTagger. Mapping is performed using minimap2 (42) with high sensitivity alignment parameters, to account for error-prone tRNA reads: `-y -ax splice -k 7 -w 3 -A 2 -B 1 -O1, 32 -E2, 0 -n 1 -m 13 -s 30 --secondary=no --MD`. The parameter file required to run *AMaNITA Preprocess* is provided in the AMaNITA GitHub repository (https://github.com/novoalab/AMaNITA/blob/master/mop/params.trna.v0.9.yaml), and it is accompanied by detailed instructions in the AMaNITA documentation, including instructions on how to run *AMaNITA Preprocess* on a demo dataset (https://github.com/novoalab/AMaNITA/wiki/Demo1), with accompanying video-guided instructions (https://github.com/novoalab/AMaNITA/wiki/Video%E2%80%90guided-AMaNITA-Preprocess).

### AMaNITA Main

AMaNITA is written in R (v4.2.1) and Python (v3.10.8). To ensure computational reproducibility and avoid installation of software dependencies, the tool executes individual processing steps within Singularity (apptainer v1.4.4-1.el9) (60) container environments. Users can launch the full pipeline via the command line, where a bash wrapper automatically manages container execution for each module.

#### Extract Metrics module

In this module, AMaNITA extracts abundance and modification information (baseQ, basecalling errors) from BAM files using of python scripts, available at the AMaNITA GitHub repository (https://github.com/novoalab/AMaNITA/tree/master/app/v0.9). First, get_counts.py returns from every BAM file the number of reads per tRNA isoacceptor (e.g., tRNA-Ala(AGC)), accumulating reads that are aligned to all its isodecoders (ie Ala-AGC-1, Ala-AGC-2, … and Ala-AGC-16). Only primary alignments in forward direction and passing predefined mapq cut-off (10) are counted. Then, get_sumerr_baseq.py aggregates for every position of each reference FASTA the following information from across 2 or more BAM files: chromosome, position, reference base, coverage and sum_err (number of mismatches at this position normalised by its coverage). The first BAM file is referred to as control, so coverage and sum_err for control are reported. For subsequent BAM files we also report coverage and sum_err, and additionally P-value and score calculated using two-sample Kolmogorov-Smirnov (KS) test (as implemented in <u>scipy.stats.ks_2samp</u>) from base qualities aggregated for a given BAM and control. Base qualities for calls that don’t match reference base are zeroed.

#### Filtering module

Filtering in AMaNITA takes place via the definition of all potential read categories based on all possible alignments of a read with regards to a reference tRNA gene (**Fig 2B**). Reads are then manipulated utilizing samtools (v1.15) (61) and bedtools (v2.30.0) (62). In this module, reads are divided into four categories: (i) full-length reads, with `start <= 1 & end >= length`, where 1 is the start and length are the start and end positions of the associated tRNA gene, respectively, (ii) 3’ capture reads, with `start > 1 & end >= length`, (iii) 5’ capture reads, with `start <= 1 & end < length`, and (iv) subset reads, with `start > 1 & end < length`. Erroneous reads and misalignments are then filtered out by following a set of three rules (**Fig 2B**): (i) All 5’ capture and subset reads are filtered out, (ii) 3’ capture reads that span less than 25bp of the tRNA gene (usually accounting to 25-35% of a tRNA gene) are filtered out, (iii) reads from any category with deletions/non-matches (denoted by N in the CIGAR string (61)) of length 10nts or larger are filtered out. This module generates filtered BAM files as output and updated TSV files from running the *<u>extract_metrics</u>* module (automatically without requiring the user to enable it) on the filtered data.

#### Technical module

Mapping, depth and coverage information are extracted with samtools (v1.15) (61) and bedtools (v2.30.0) (62). The module computes the following metrics, and the obtained results are plotted into the report: i) ligation efficiency score; and ii) degradation score. The ligation efficiency score informs on the efficiency of adapter ligation by assessing the proportion of reads lacking ∼15nts at the 5’ end of RNA molecules, and is calculated as follows:

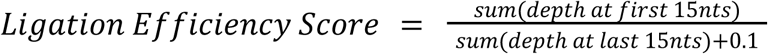

The degradation score informs on potential RNA degradation, utilizing the relative proportion of full-length and 3’ capture reads, and is defined as follows:

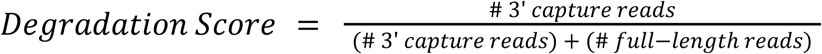

If the *filtering* module is enabled, the *technical* module is run both before (“pre”) filtering and after (“post”) filtering, in order to supply quality control and technical information on the data obtained at both stages.

#### Batch module

Batch effect investigation and correction in AMaNITA is carried out in three steps, following best practices set in the field of bioinformatics for Illumina data and adjusting them for DRS and tRNA data (47). In the first step, we assess the possible sources of technical variation (batch effects) through exploratory data analysis and visualization. Our approach includes the assessment of the connectivity and balance of the experimental design in order to inform the user of potential caveats in the experimental design that would inhibit batch effect adjustment and proper statistical analyses downstream. In addition, we perform Principal Component Analysis (PCA) and hierarchical clustering, separately on the abundance and modification data.

In the second step, we assess the effect of suspected or unknown sources of technical variation through the use of rigorous statistical frameworks. To assess suspected batch effects, defined by the user, we perform Principal Variance Component Analysis (PVCA) adjusted for tRNAs (vsn v3.66.0, lme4 v1.1-30) (44), which allows us to estimate the variance explained by a specific batch effect. To assess unknown sources of technical variation by harnessing the data itself, we perform Surrogate Variable Analysis (SVA) (v3.46.0) (45, 46), which allows us to agnostically identify Surrogate Variables (SVs) removing the need to know important batch effects *a priori*. We further assess the association of the significant batch effects and surrogate variables identified to the Principal Components in order to obtain the full picture of their effect.

In the third step, we use a pre-determined set of rules for selecting the variables that should be accounted and corrected for: (i) *Criterion 1*: if a suspected batch effect (defined by the user) accounts for more than 10% of the total variance of the data, it should be included in the model, and (ii) *Criterion 2*: all identified surrogate variables (SVs) that are not confounded with the biological group of interest or suspected batch effects (r < 0.8, so as to avoid collinearity) are included in the model; however, for the cases where SVs are correlated to the biological group of interest (r > 0.4), AMaNITA gives a warning as it anticipates the undesired removal of biological signal along with the technical noise, and prompts the user to further investigate whether, conversely, the SV is robust enough to remove only technical noise. Upon identification of those important sources of technical variation, the *<u>batch</u>* module performs batch effect correction for visualization purposes only by utilizing limma’s `removeBatchEffect` function (v3.54.2) (63) to obtain batch-corrected data, run PCA on it, and visualize. In addition, it automatically informs the *<u>pairwise</u>* module of batch effects to account for in the linear model, while providing the same information to the user. We should note, however, that it falls to the user’s discretion to confirm that a model is appropriate, and especially to confirm that the balance of experimental design is not violated. To do so, AMaNITA provides all the required plots and tables to aid the user in choosing the appropriate model.

#### Pairwise module

Pairwise comparisons are carried out by AMaNITA between two groups of interest, specified in the user-supplied metadata file. Differential abundance analysis is carried out with DESeq2 (v1.38.3) (48), after confirming that the tRNA data follows the expected negative binomial distribution. Differential modification analysis by utilizing basecalling errors is carried out with edgeR in methylation mode (v3.40.2) (51, 52). Informed by the *<u>batch</u>*module, the *<u>pairwise</u>* module can automatically incorporate important technical covariates (suspected batch effects or identified surrogate variables) into the linear model, allowing for proper batch effect handling. Differential modification analysis by utilizing the baseQ information is carried out with Kolmogorov-Smirnov test by comparing the baseQ distributions between two groups of interest. We also offer the results of the ΔSumErr differential modification analysis for groups with multiple replicates by defining ΔSumErr as follows:

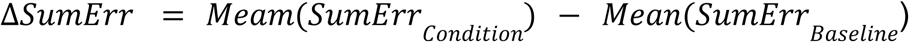

#### Pairwise Custom

This module performs the same processes as the *<u>pairwise</u>*module, but allows for additional flexibility in defining the linear model to be utilized for the differential analyses and in expanding group comparisons to <u>1 vs many</u> or <u>many vs many</u>. In the first case, the user can supply additional covariates of interest to run a more complex model; for example, while the *<u>batch</u>*-informed *<u>pairwise</u>* module would run `∼ condition + batch`, *<u>pairwise_custom</u>* allows for designs like ∼ `condition + batch + sex + age`. In the second case, AMaNITA utilizes the capability of the `DESeq::results` and `edgeR::makeContrasts` function to define contrasts with more that 1 groups; for example, it is possible to run the following: `res <-results(dds, contrast = list(c(“condA”, “condB”), “control”), listValues = c(1/2, -1))`, allowing for more complex comparisons between multiple groups.

### AMaNITA Reports and Plotting

HTML reporting in AMaNITA is inspired by SQANTI3 (64), and it utilizes plotly (v4.10.0) (65), pander (v0.6.5), htmltools (v0.5.3), and DT (v0.26) to render the final report. Further plotting utilizes ggplot2 (v3.3.6), ggridges (v0.5.4), ggplotify (v0.1.0), ggfortify (v0.4.14), ggrepel (v0.9.1), and ggridges (v0.5.4) for general plots; igraph (v1.3.5) and ggraph (v2.1.0) for graph plots; ggdendro (v0.1.23) for dendrograms; corrplot (v0.92) for correlation plots; ComplexHeatmap (v2.14.0) for heatmaps; EnhancedVolcano (v1.16.0) for volcano plots. Data handling is done via the use of tidyr (v1.2.1), stringr (v1.4.1), magrittr (v2.0.3), data.table (v1.14.4), and dplyr (v1.0.10).

### Annotations used in this work

Reference sequences for mature *S. cerevisiae* tRNAs were retrieved from GtRNAdb2 (66). GtRNAdb2 reports 275 tRNA sequences annotated in the *S. cerevisiae* genome. Only sequences that were at least 5% divergent at nucleotide level (including ligated 5’ and 3’ oligos) were kept in the final reference, which retained one reference tRNA gene per tRNA isoacceptor. The yeast tRNA reference file used in this work is available in GitHub (https://github.com/novoalab/Nano-tRNAseq and https://github.com/novoalab/AMaNITA/blob/master/aux/ref_v0.9/sacCer3.fa.bed). Modifications for yeast tRNAs were obtained from MODOMICS (67) and the canonical position was manually curated using published literature, as previously described (20).

## CODE AVAILABILITY

AMaNITA is publicly available on GitHub (https://github.com/novoalab/AMaNITA), and is also provided as a supplementary ZIP file accompanying this manuscript. A stable Zenodo DOI will be generated once the code has been peer-reviewed. AMaNITA is written in bash, R and Python, and individual modules are executed within a Singularity container environment. Users can launch the full pipeline via the command line, where the wrapper automatically manages container execution for each internal submodule.

## ACKNOWLEDGEMENTS

We thank all the members of the Novoa lab for their insightful discussions and for beta-testing AMaNITA. We also thank Sonia Tarazona and Carolina Monzó for the insightful discussions on statistical analyses. This project has received funding from the European Union’s Horizon 2020 research and innovation programme under the Marie Sklodowska-Curie grant agreement No 101072892 (Marie Sklodowska-Curie Actions Doctoral Network LongTREC). EMN and this project received funding from the project PID2024-157315NB-I00; the European Union’s Horizon Europe through the European Research Council under the grant agreement number 101042103 and 101187456. Views and opinions expressed are however those of the author(s) only and do not necessarily reflect those of the European Union. Neither the European Union nor the granting authority can be held responsible for them. We acknowledge support of the Spanish Ministry of Science and Innovation through the Centro de Excelencia Severo Ochoa (CEX2020-001049-S, MCIN/AEI /10.13039/501100011033), and the Generalitat de Catalunya through the CERCA programme and to the EMBL partnership. We are grateful to the CRG Core Technologies Programme for their support and assistance in this work.

## AUTHOR CONTRIBUTIONS

XLK wrote AMaNITA, benchmarked the software and performed all bioinformatic data analyses included in this work. LPP wrote code for automatization of Nano-tRNAseq data basecalling, mapping and tRNA quantification steps, and built tRNA mapping references. LLL cultured yeast, prepared and sequenced Nano-tRNAseq libraries. AO performed the mouse work, including mice husbandry, dissection, RNA extraction and Nano-tRNAseq libraries. LC and JP implemented the code for basecalling, mapping and quantifying tRNAs into a NextFlow workflow (AMaNITA Preprocess). EMN conceived and supervised the work. XLK built the figures and drew the AMaNITA logo. XLK and EMN discussed and interpreted the results. XLK and EMN wrote the manuscript, with contributions from all authors.

