## Supplementary Figures for "AMaNITA: an end-to-end workflow for native tRNA nanopore sequencing data analysis"

**Figure S1. Flowchart of AMaNITA software.** This flowchart diagram representation schematically outlines the major entry points, processes, and output which make up AMaNITA. Starting at entry point A (POD5 files), the user can employ *AMaNITA Preprocess* to carry out basecalling, demultiplexing, and alignment, and finally obtain aligned BAM files. Starting at entry point B (BAM files), the user does not have to run AMaNITA Preprocess and can directly employ AMaNITA main and all of its available modules, including *extract_metrics* to obtain abundance and modification TSV files necessary for some of the analyses, *filtering* to remove spurious reads and erroneous alignments, *technical* to obtain quality control information on the dataset as a whole, *batch* to assess the effect of unwanted variables and their contribution to the data in the form of technical noise, *pairwise* to perform pairwise (between 2 groups) differential abundance and differential modification analyses, and *pairwise_custom* that further expands the flexibility of the pairwise module. AMaNITA can also be employed at entry point C (TSV) files, but the modules that can be run in this case are limited to *batch*, *pairwise*, and *pairwise_custom*. The results from all modules comprising *AMaNITA Main* are collapsed into the AMaNITA Report, and are also provided in the form of figures and accompanying tables.

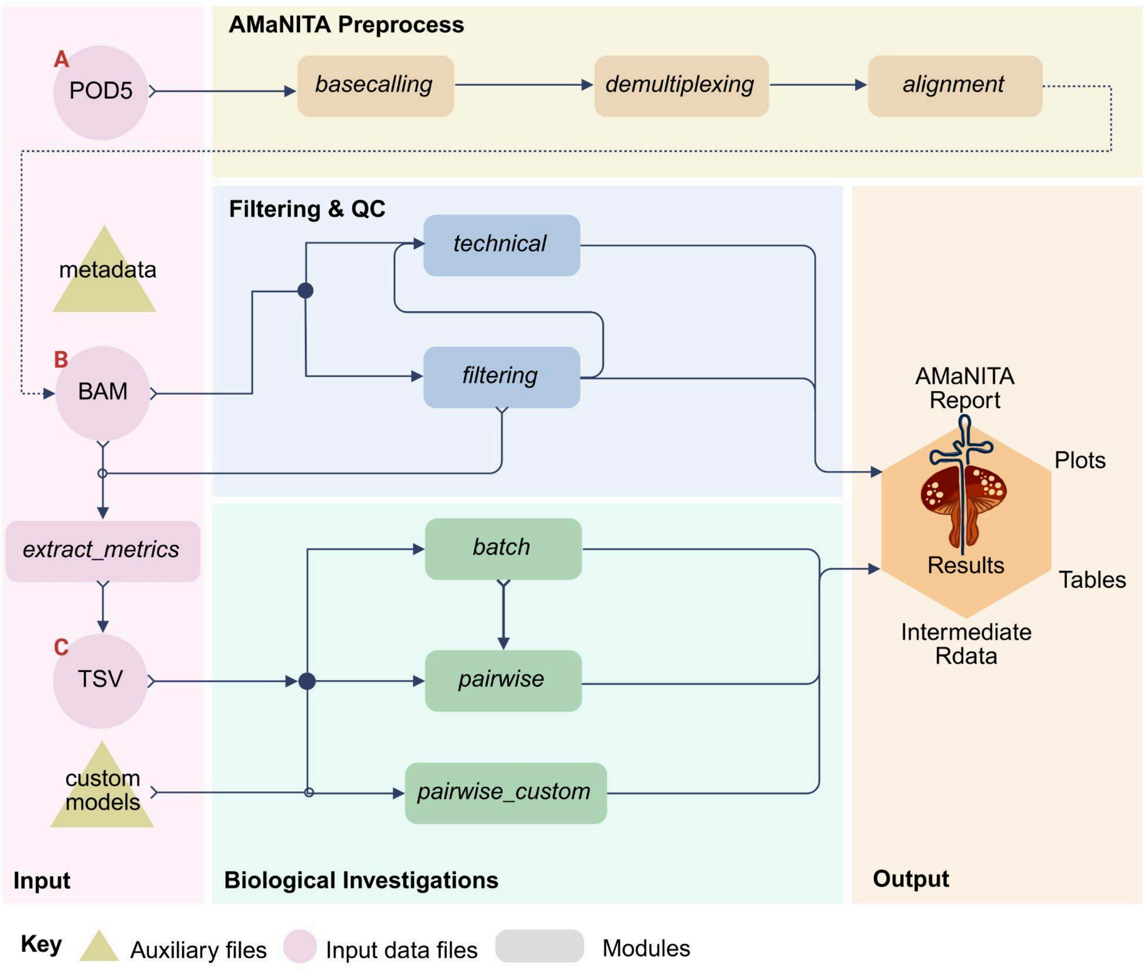

**Figure S2. The His-GTG case: Identifying improper adapter ligation through AMaNITA's *technical* module. (A)** Per sample coverage percentage distribution of reads mapping to His-GTG, demonstrating the higher percentage of truncated reads (coverage < 100%), with small peaks at ~50% and ~80% showing the existence of reads which cover half or ~4/5 of the tRNA gene instead of the total gene (full-length); a consistent observation for this tRNA due to improper adapter ligation. **(B)** Per tRNA Ligation Efficiency score barplot, demonstrating how His-GTG is a consistent outlier and, therefore, setting an internal baseline for ligation efficiency. **(C)** Per tRNA Degradation score barplot, demonstrating how His-GTG is a consistent outlier and, therefore, setting an internal baseline for degradation, even if the reason for this behaviour is improper adapter ligation.

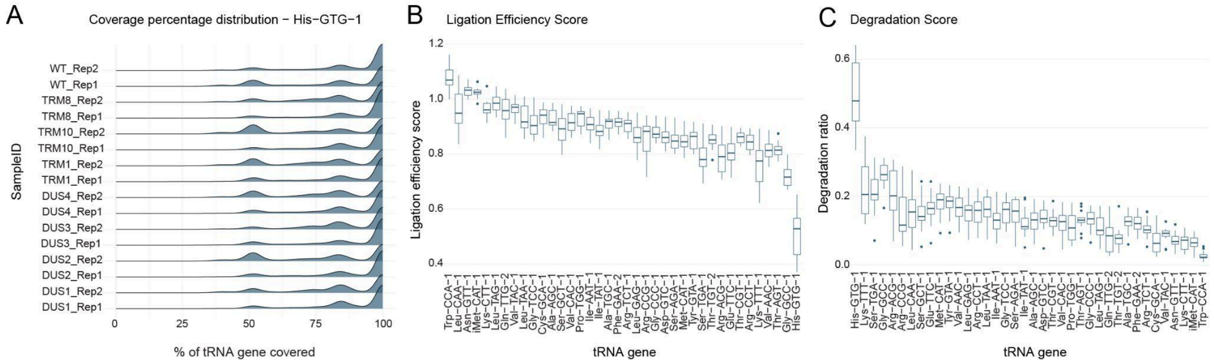

**Figure S3. Identification of sources of technical variation and considerations for selecting the appropriate linear model. (A)** Hierarchical clustering of the abundance (left) and modifications data (right), coloured by biological group and suspected batch effect reveals a persistent source of technical variation. **(B)** Applying AMaNITA's criteria for selecting the most appropriate linear model for the differential abundance and differential modifications analyses (left), taking into account experimental design connectivity and balance (right).

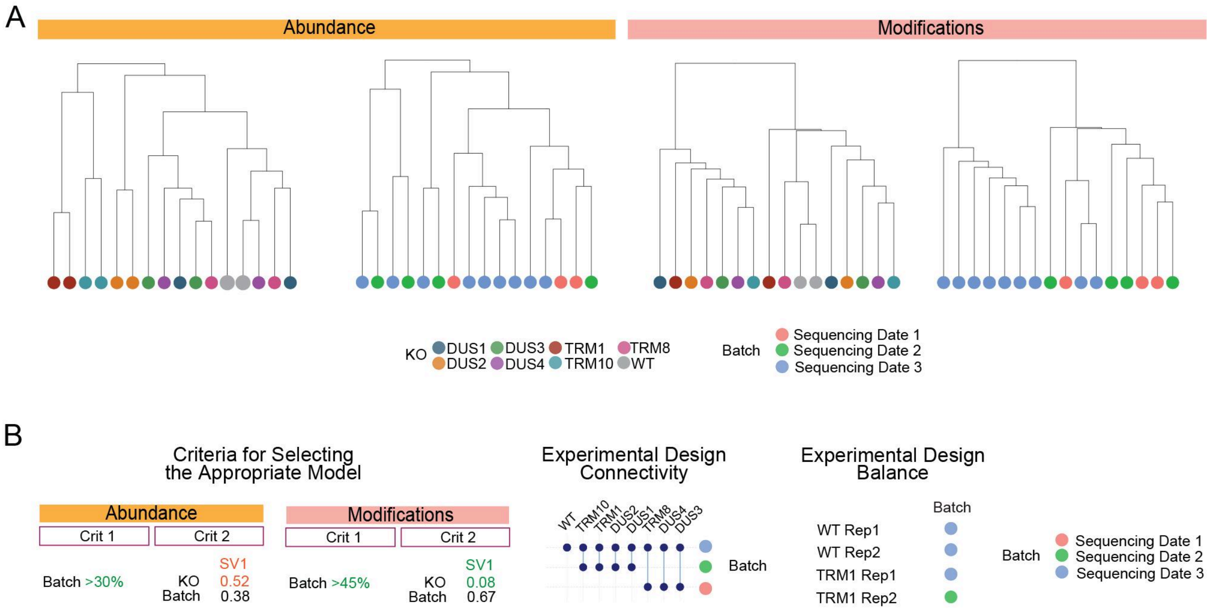

**Figure S4. Screenshot from the AMaNITA Report.** The screenshot was taken from the Batch Effect Investigation section depicting the table of suggested linear models and the accompanying warnings with regards to surrogate variable correlation with the group of interest and experimental design balance, for the abundance data.

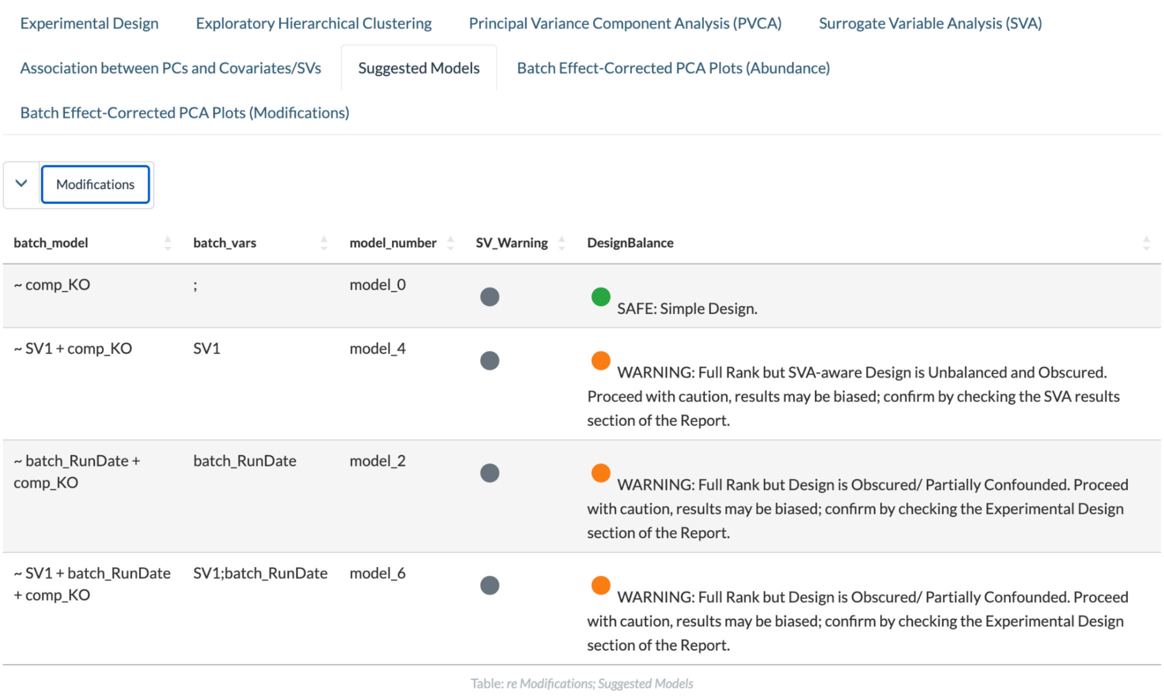


| batch_model | batch_vars | model_number | SV_Warning | DesignBalance |
|---|---|---|---|---|
| ~ comp_KO | ; | model_0 | | SAFE: Simple Design. |
| ~ SV1 + comp_KO | SV1 | model_4 | | WARNING: Full Rank but SVA-aware Design is Unbalanced and Obscured. Proceed with caution, results may be biased; confirm by checking the SVA results section of the Report. |
| ~ batch_RunDate + comp_KO | batch_RunDate | model_2 | | WARNING: Full Rank but Design is Obscured/ Partially Confounded. Proceed with caution, results may be biased; confirm by checking the Experimental Design section of the Report. |
| ~ SV1 + batch_RunDate + comp_KO | SV1;batch_RunDate | model_6 | | WARNING: Full Rank but Design is Obscured/ Partially Confounded. Proceed with caution, results may be biased; confirm by checking the Experimental Design section of the Report. |

Table: *re Modifications; Suggested Models*

**Figure S5. Differential abundance analysis results for the TRM1 KO vs WT comparison.** Volcano plot (left) and heatmap (right) generated by AMaNITA.

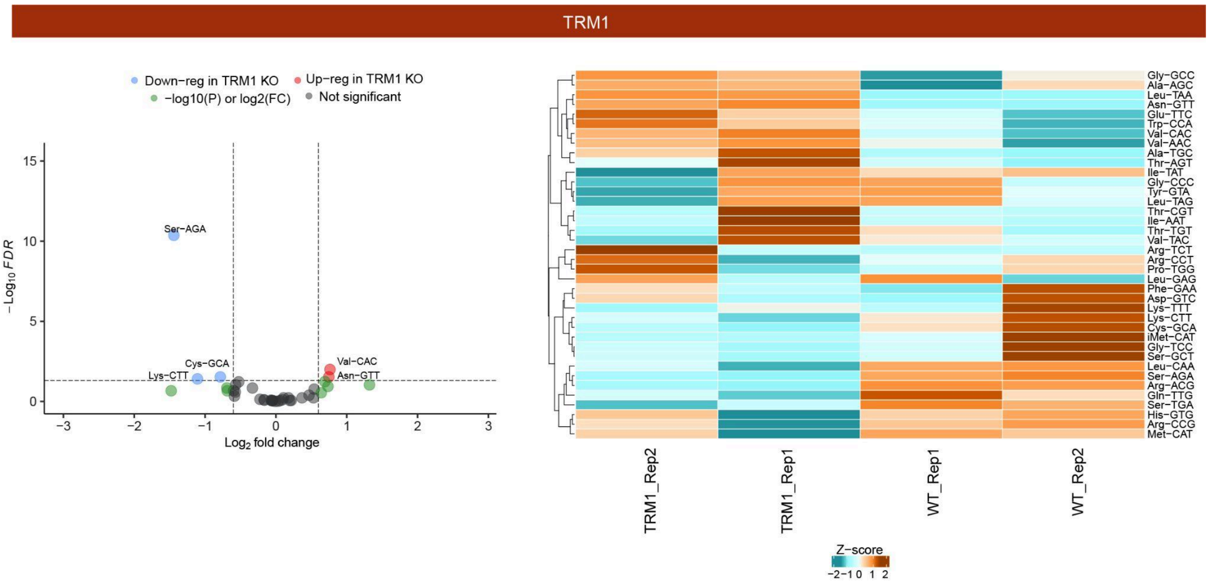

**Figure S6. Differential abundance analysis results for the Dnmt2 KO vs WT comparison (RNA002). (A)** Replicability scatterplots of WT (left) and KO (right) replicates, coloured by tRNA, as generated by AMaNITA. **(B)** Volcano plot generated by AMaNITA when comparing Dnmt2 and WT datasets. **(C)** Heatmap generated by AMaNITA, depicting the tRNA abundance across the 4 samples, with Z-scored values.

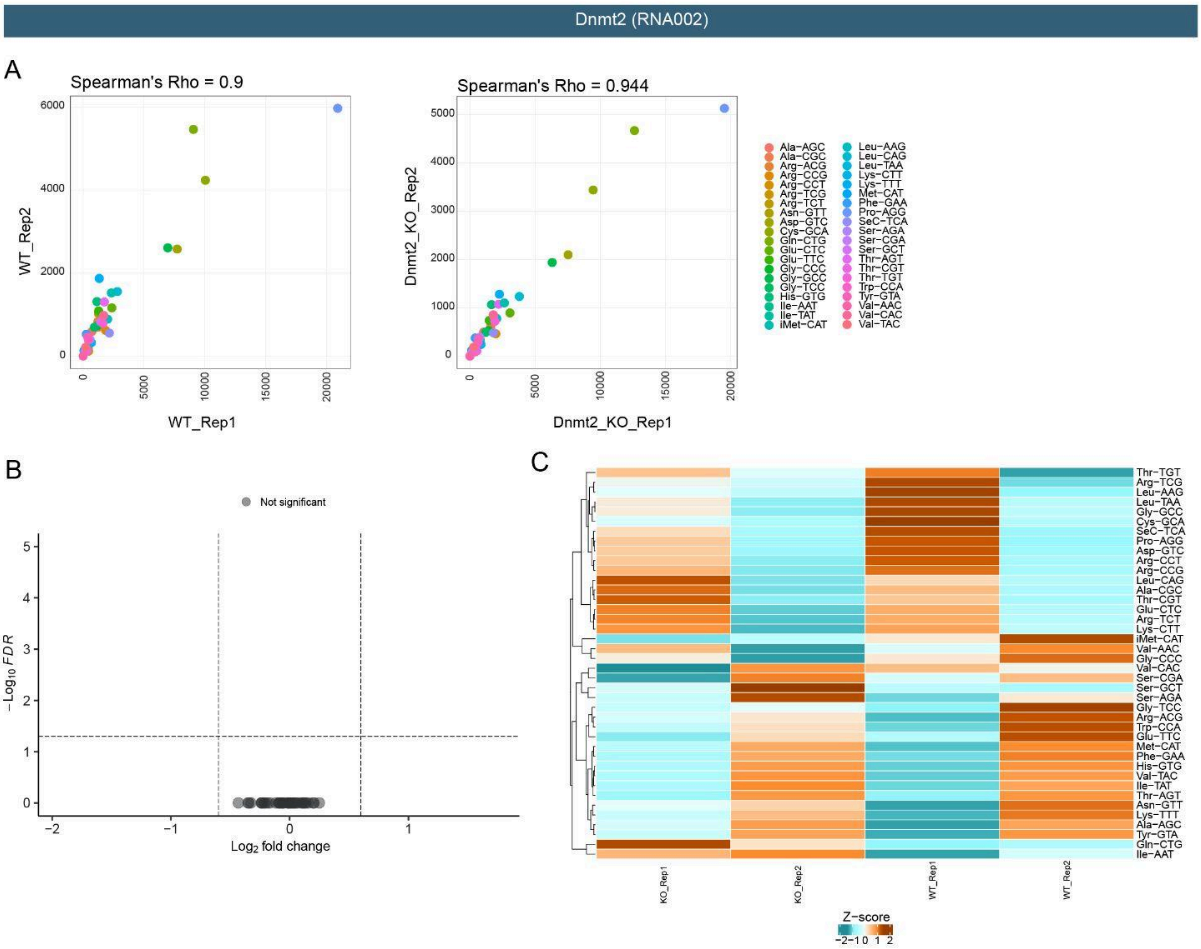

**Figure S7. Differential modification results for the TRM1, DUS1, DUS3, and DUS4 KO strains versus the wild type strain.** Positional heatmaps with the differential modifications analysis results in three different cases: on a dataset with a dominant batch effect and an unbalanced experimental design, adjusting for batch effect in the aforementioned unbalanced dataset, and on a balanced dataset and/or not affected by batch effect. tRNAs are marked in **bold** and coloured if they harbor a modification at the expected position(s) according to the *Modomics* annotation.

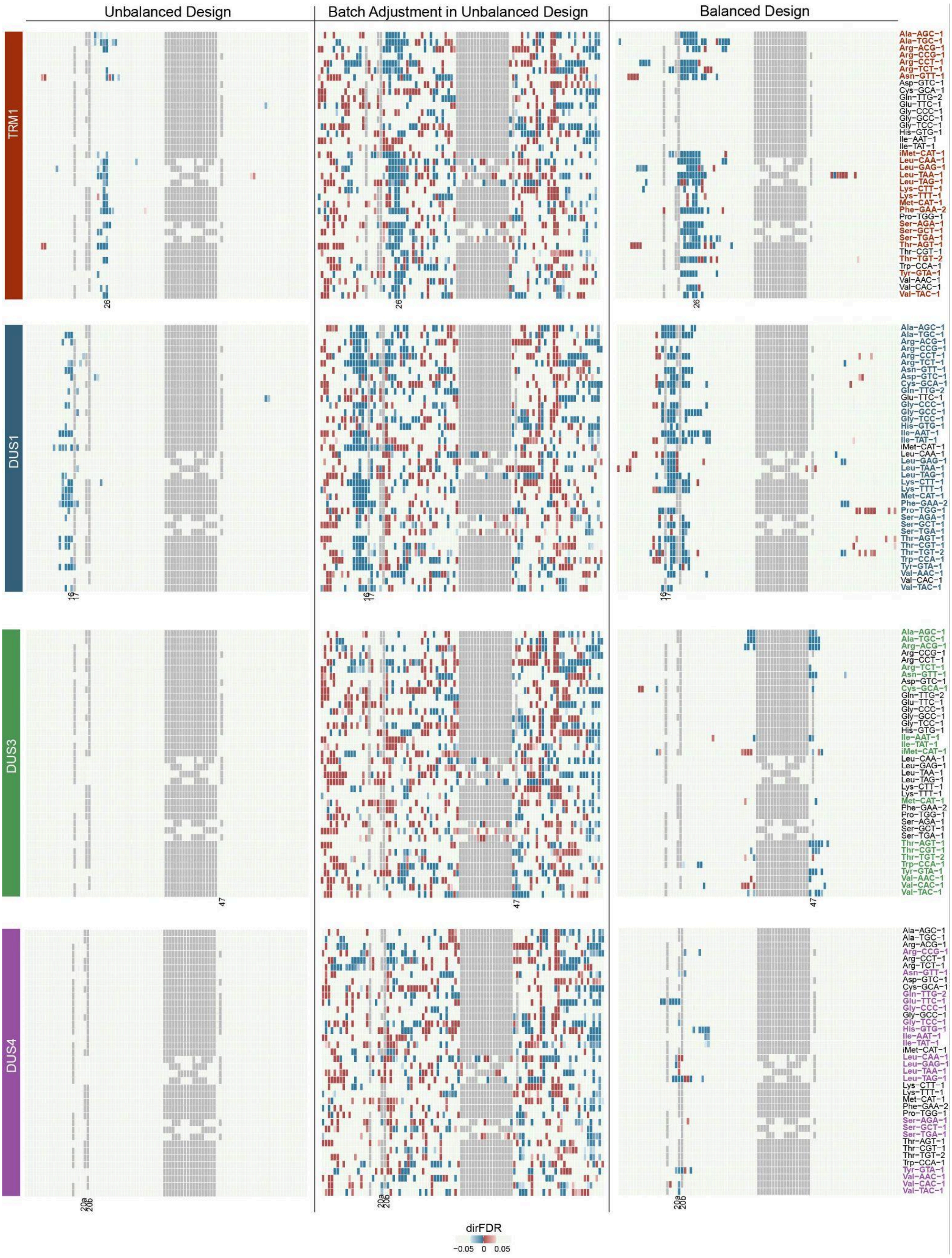
