## Supplemental File 1 for "AMaNITA: an end-to-end workflow for native tRNA nanopore sequencing data analysis": FileS1.html

AMaNITA Report


### AMaNITA Report

###### 28 May, 2026


---


### Summary

#### Settings

#### Metadata Table

#### Filter Details

#### Reference Assumptions

#### Generated Plots

##### Filtering Module

Plots generated by the filtering module can be found at: `/users/enovoa/lkatopodi/Projects/appv0.9/dnmt2_RNA002/cyto/ftbp/filtering/plots`

##### Technical Module

Plots generated by the technical module (pre-filtering) can be found at: `/users/enovoa/lkatopodi/Projects/appv0.9/dnmt2_RNA002/cyto/ftbp/technical_{pre or post}/plots`

##### Batch Module

Plots generated by the batch module can be found at: `/users/enovoa/lkatopodi/Projects/appv0.9/dnmt2_RNA002/cyto/ftbp/batch/{comp_variable}/plots`

##### Pairwise Module

Plots generated by the pairwise module can be found at: `/users/enovoa/lkatopodi/Projects/appv0.9/dnmt2_RNA002/cyto/ftbp/pairwise/{comp_variable}/{comp_ID}/plots`

##### Report

Plots generated for the report can be found at: `/users/enovoa/lkatopodi/Projects/appv0.9/dnmt2_RNA002/cyto/ftbp/report/plots`

Input abundance and modification data tables can be found at: `/users/enovoa/lkatopodi/Projects/appv0.9/dnmt2_RNA002/cyto/ftbp/report`

### Quality Control Metrics

#### Input Abundance Data

##### Table

##### Total Reads

png 2 png 2

#### Exploratory PCAs

##### Abundance

###### comp\_KO

###### batch\_RunID

##### Modifications

###### comp\_KO

###### batch\_RunID

#### Filtered Reads

##### Total Reads

##### Filtering Stats

##### Number of Reads filtered

##### Normalized Number of Reads filtered

##### Percentage of Reads filtered

```
       used  (Mb) gc trigger  (Mb) max used  (Mb)
```

Ncells 8457294 451.7 15610938 833.8 12309565 657.5 Vcells 15541565 118.6 26336561 201.0 26336561 201.0

#### tRNA Fractions

##### Isoacceptor

###### Pre-filtering

###### Post-filtering

##### tRNA Gene

###### Pre-filtering

###### Post-filtering

##### Normalized per Isoacceptor

###### Pre-filtering

###### Post-filtering

#### Read Composition per Sample

##### Overview

###### Pre-filtering

###### Post-filtering

##### Full-length Reads

###### Pre-filtering

###### Post-filtering

##### 3’ Reads

###### Pre-filtering

###### Post-filtering

##### 5’ Reads

###### Pre-filtering

##### Subset Reads

###### Pre-filtering

#### Read Composition per Sample

##### Overview

###### Pre-filtering

###### Post-filtering

##### 3’ Reads

###### Pre-filtering

###### Post-filtering

##### 5’ Reads

###### Pre-filtering

##### Subset Reads

###### Pre-filtering

#### Ligation Efficiency Score

##### Pre-filtering

##### Post-filtering

#### Degradation Score

##### Pre-filtering

##### Post-filtering

```
       used  (Mb) gc trigger  (Mb) max used  (Mb)
```

Ncells 8472935 452.6 15610938 833.8 15610938 833.8 Vcells 15597319 119.0 51506801 393.0 38419197 293.2

### Batch Effect Investigation

#### comp\_KO

##### Experimental Design

###### Experimental Design Connectivity Rationale

###### Sewing Plot

Overlapping horizontal lines = Connected Design

Vertical gaps = Disconnected Design

Dots without Connection = Isolated Islands (Obscured)

###### batch\_RunID

###### Connectivity Graph

###### batch\_RunID

###### Experimental Design Balance

##### Exploratory Hierarchical Clustering

###### Abundance

###### comp\_KO

###### batch\_RunID

###### Modifications

###### comp\_KO

###### batch\_RunID

##### Principal Variance Component Analysis (PVCA)

###### Abundance

###### Potential Batch Effects

###### All Effects

PVCA error: Error: number of levels of each grouping factor must be < number of observations (problems: comp\_KO:batch\_RunID)

Showing PVCA results with no Interaction Terms

###### Batch Variables to Adjust for

###### Other Effects

###### Modifications

###### Potential Batch Effects

###### All Effects

PVCA error: Error: number of levels of each grouping factor must be < number of observations (problems: comp\_KO:batch\_RunID)

Showing PVCA results with no Interaction Terms

###### Batch Variables to Adjust for

###### Other Effects

##### Surrogate Variable Analysis (SVA)

###### Abundance

###### Metadata

##### SV1

###### Modifications

###### Metadata

##### SV1

##### Association between PCs and Covariates/SVs

###### Abundance

###### Correlation w/ PCs

###### Correlation w/ SVs

###### Modifications

###### Correlation w/ PCs

###### Correlation w/ SVs

##### Suggested Models

###### Abundance

###### Modifications

##### Batch Effect-Corrected PCA Plots (Abundance)

###### model\_2

###### comp\_KO

###### batch\_RunID

###### PCA Loadings Plot

##### Batch Effect-Corrected PCA Plots (Modifications)

###### model\_2

###### comp\_KO

###### batch\_RunID

###### PCA Loadings Plot

```
       used  (Mb) gc trigger  (Mb) max used  (Mb)
```

Ncells 8536263 455.9 15610938 833.8 15610938 833.8 Vcells 16106070 122.9 51506801 393.0 38419197 293.2

### dnmt2\_KO vs WT

Below are the results for the comparison: comp\_KO.dnmt2\_KO\_vs\_WT

#### Differential Abundance

##### Examined Models

##### DESeq2 Results Table

###### model\_0

###### model\_2

##### Volcano Plots

###### model\_0

###### p-value

###### FDR

###### model\_2

###### p-value

###### FDR

##### Heatmaps

###### model\_0

###### Normalized Abundance

###### Normalized per AA

###### model\_2

###### Normalized Abundance

###### Normalized per AA

#### Differential Modifications

##### Examined Models

##### Results Tables

###### edgeR on basecalling errors

###### model\_0

###### model\_2

###### baseQ KS test

##### Volcano Plots

###### edgeR - model\_0

###### p-value

###### FDR

###### edgeR - model\_2

###### p-value

###### FDR

###### baseQ KS test

###### p-value

###### FDR

##### Positional Heatmaps

###### edgeR - model\_0

###### FDR

###### FDR (no logFC cutoff)

###### logFC

###### Heuristic

###### edgeR - model\_2

###### FDR

###### FDR (no logFC cutoff)

###### logFC

###### Heuristic

###### baseQ KS test

###### FDR

###### FDR (no KS score cutoff)

###### KS score

###### DSumErr

##### Positional Barplots

###### edgeR - model\_0

###### edgeR - model\_2

###### baseQ KS test

```
       used  (Mb) gc trigger  (Mb) max used  (Mb)
```

Ncells 8531162 455.7 15610938 833.8 15610938 833.8 Vcells 17933428 136.9 51506801 393.0 51506801 393.0
